# Intracellular absorption underlies collective bacterial tolerance towards an antimicrobial peptide

**DOI:** 10.1101/314138

**Authors:** Fan Wu, Cheemeng Tan

## Abstract

The collective tolerance towards antimicrobial peptides (APs) is thought to occur primarily through mechanisms associated with live bacterial cells. In contrast to the focus on live cells, we discover that the LL37 antimicrobial peptide kills *Escherichia coli*, forming a subpopulation of dead cells that absorbs the remaining LL37 into its intracellular space. Combining mathematical modeling with population and single-cell experiments, we show that bacteria absorb LL37 at a timing that coincides with the permeabilization of their cytoplasmic membranes. Furthermore, we show that one bacterial strain can absorb LL37 and protect another strain from killing by LL37. Finally, we demonstrate that the intracellular absorption of LL37 can be reduced using a peptide adjuvant. In contrast to the existing collective tolerance mechanisms, we show that the dead-bacterial absorption of APs is a dynamic process that leads to emergent population behavior, and the work suggests new directions to enhance the efficacy of APs.

## Introduction

Antimicrobial peptides (APs) are small peptides (normally less than 10kDa) that counter bacterial pathogens in host innate immune systems (Cole & Nizet, 2016; Nizet et al., 2001) and are being developed as new sources of antibacterial agents (Gordon, Romanowski, & McDermott, 2005; Hancock & Sahl, 2006). Major efforts in the field have focused on interaction dynamics between APs and bacterial components. For instance, after the initial contact to bacterial membranes driven by the cationic domain of APs, the hydrophobic domain of APs facilitates the insertion of APs into lipid bilayers, leading to membrane permeabilization and cell death (Nguyen, Haney, & Vogel, 2011; Teixeira, Feio, & Bastos, 2012). Previous studies have also shown that some APs can target DNA (Hsu et al., 2005; Podda et al., 2006) and intracellular proteins (Kragol et al., 2001; Otvos, Snyder, Condie, Bulet, & Wade, 2005). However, beyond the direct interaction between APs and bacterial targets, APs can be tolerated by certain bacterial species through collective mechanisms. The collective tolerance mechanisms are relatively well-studied for classical antibiotics (Meredith, Srimani, Lee, Lopatkin, & You, 2015; Vega & Gore, 2014) when compared to antimicrobial peptides, and their implication on antibiotic treatment is well demonstrated in the literature (Chait, Palmer, Yelin, & Kishony, 2016; Hol, Hubert, Dekker, & Keymer, 2016; Vega, Allison, Samuels, Klempner, & Collins, 2013; Yurtsev, Chao, Datta, Artemova, & Gore, 2013). For antimicrobial peptides, bacteria may exhibit collective tolerance through mechanisms such as membrane-displayed proteases that degrade APs (Johansson et al., 2008; Schmidtchen, Frick, Andersson, Tapper, & Bjorck, 2002; Sieprawska-Lupa et al., 2004) and secreted molecules including lipids, vesicles, and proteins that titrate APs (Campos et al., 2004; Cole et al., 2010; Frick, Åkesson, Rasmussen, Schmidtchen, & Björck, 2003; Spinosa et al., 2007). The titration mechanism occurs due to electrostatic interactions between cationic APs and negatively-charged molecules or surfaces (Bucki, Byfield, & Janmey, 2007; Frick et al., 2003; Llobet, Tomas, & Bengoechea, 2008; Starr, He, & Wimley, 2016; Weiner, Bucki, & Janmey, 2003).

To understand collective tolerance caused by the titration mechanism, it is necessary to first track the localization and distribution of APs in a bacterial population. However, previous results have been contradictory because the minimum inhibitory concentration (MIC) of APs is at least two-log folds higher than the amount necessary to kill a single bacterium, suggesting an unknown titration source. Through fluorescence spectroscopy of the AP PMAP-23, it is found that bacteria are killed when the AP molecules saturate the total surface area of bacterial membranes with 10^6^-10^7^ peptides per cell (Roversi et al., 2014). Instead, another study has shown that MIC of AP Pexiganan requires ~10^9^ peptides per cell (Jepson, Schwarz-Linek, Ryan, Ryadnov, & Poon, 2016), which is much higher than the necessary amount of AP to saturate the surface area of a single bacterium (Jepson et al., 2016; Wimley, 2010). In addition, if the membrane of live cells is the only titration source of APs, MIC of bacteria must increase linearly with the inoculum size. This expectation has also been proven wrong in the literature (Jepson et al., 2016). What is the hidden factor that contributes significantly to the titration of AP in a bacterial population that does not exhibit any of the known tolerance mechanisms (i.e., lipid shedding and protease display)? Answers to the question may lead to a new explanation of collective tolerance dynamics during AP treatment and innovative methods to enhance the efficacy of APs.

Instead of focusing on live bacterial cells following current thoughts in the field, we find that dead bacterial cells can serve as a major titration source of an AP. We discover that LL37, which is a cathelicidin family AP from human, permeabilizes cytoplasmic membranes of a subpopulation of bacteria (*Escherichia coli*), which then absorbs LL37 into its intracellular space. The titration of LL37 by permeabilized bacteria forms a negative feedback response to LL37 treatment, generating emergent collective tolerance dynamics that cannot be predicted without the AP-absorption mechanism. Specifically, we track the dynamics of LL37 in bacterial populations using both single-cell and population measurements based on previous work (Fantner, Barbero, Gray, & Belcher, 2010; Sochacki, Barns, Bucki, & Weisshaar, 2011). We first rule out existing AP-tolerance mechanisms in our model system, including the modification of bacterial surface charge (Fabretti et al., 2006; Guo et al., 1998; Kovacs et al., 2006; Poyart et al., 2003; Starner, Swords, Apicella, & McCray, 2002), the inactivation of APs by surface shielding (Campos et al., 2004; Cole et al., 2010; Spinosa et al., 2007), and the cleavage of LL37 (Johansson et al., 2008; Schmidtchen et al., 2002; Sieprawska-Lupa et al., 2004). Next, we show that the amount of free LL37 in the bacterial culture is reduced through bacterial absorption, which allows a subpopulation of *E. coli* to grow and repopulate the culture. We also present single-cell data and perturbation experiments that confirm the AP-absorption mechanism.

Furthermore, we demonstrate that the AP-absorption leads to emergent cross-bacterial-strain protection against LL37. To illustrate the importance of understanding the AP-absorption mechanism, we show that a peptide adjuvant can be supplemented to reduce the absorption of APs. The AP-absorption mechanism may be generalizable to other bacterial species and APs, and it may be considered in the new design of AP-treatment that enhances the efficacy of APs.

## Results

### Bacterial population recovers from initial killing by LL37 through a non-heritable mechanism

To establish the experimental conditions for our study, we first investigate real-time growth dynamics of *E. coli* under LL37 treatment. The *E. coli* BL21PRO strain expresses *lux* genes (BP-lux), leading to luminescence that is tracked as a surrogate of bacterial viability using a platereader (Yeh, Tschumi, & Kishony, 2006). Luminescence intensity is widely used to report bacterial metabolic state under antimicrobial treatment because it exhibits higher sensitivity and larger dynamic range than optical density (Bjarnason, Southward, & Surette, 2003; Kishony & Leibler, 2003; Yeh et al., 2006). For this experiment, we initiate cultures using the M9 medium with ~10^3^-10^4^ CFU/μl of bacteria (See pre-growth protocol 1 in Methods Section M1) and measure their growth dynamics in a 96-well plate supplemented with 2-fold dilutions of LL37 using a platereader for at least 14 hours (See Methods Section M2). In typical antibiotic tests using batch cultures, bacteria would either grow or be inhibited by the antibiotic for at least 24 hours before the emergence of resistant mutants (Tan et al., 2012). Instead, for LL37, we find that bacterial populations are inhibited (decline in luminescence intensity) by LL37 at 6.75μg/ml (Fig. 1a, black dash line) in the first six to eight hours, after which they re-grow at the same rate as the untreated bacteria (Fig. 1a, black line). We confirm this trend using CFU of the bacteria (Fig. S1). The recovery dynamic does not occur with LL37 at 13.5μg/ml within the experimental duration (Fig. 1a, grey line). To validate the mode of actions of LL37, we treat wild-type BL21PRO (WT-BP) with 13.5μg/ml LL37 for 2 hours. We find strong signals of phosphatidylserine (PS) exposure (Fig. 1c and Fig. S2a) and propidium iodide (PI) staining (Fig. 1d and Fig. S2b), which have been used as markers for bactericidal antibiotics (Dwyer, Camacho, Kohanski, Callura, & Collins, 2012), and cell permeabilization and death (Davey & Hexley, 2011) in previous studies. Unless otherwise noted, we use the intermediate concentrations (i.e., sub-MIC) of LL37 that allow bacterial recovery to reveal the unknown collective tolerance mechanism. The use of sub-MIC concentrations is well accepted in the study of tolerance mechanisms (Müller et al., 2016; Pader et al., 2017; Tan et al., 2012) because bacterial growth dynamics are sensitive to tolerance mechanisms at sub-MIC, increasing the feasibility of detecting the tolerance mechanisms. Despite the use of sub-MIC for our study, the revealed collective tolerance mechanisms will occur at AP concentrations above MIC and potentially reduce the efficacy of the AP.

**Figure 1.**
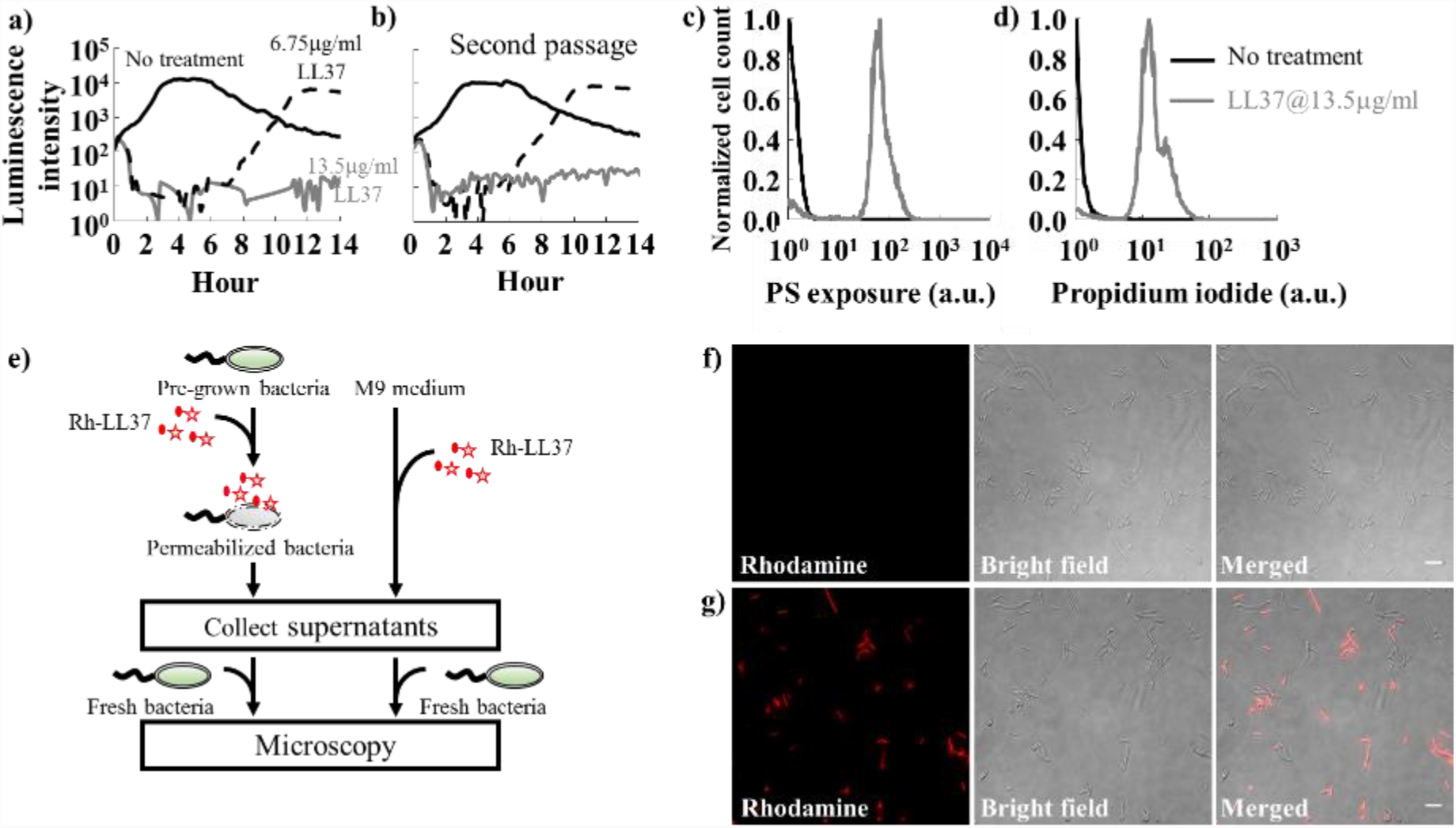
Studying collective tolerance mechanisms using LL37 and *Escherichia coli*. **a)** Population dynamics of *E. coli* are tracked using their luminescence intensity. Bacteria treated with LL37 at two concentrations (black dash line and grey line) demonstrate initial killing (before 6 hours) compared to the one without treatment (black line). However, the bacterial population treated with 6.75μg/ml of LL37 recovers after around 8 hours. See Methods Section M2. **b)** Recovered bacteria from LL37 treatment at 6.75μg/ml (Fig. 1a black dash line) is collected and grown overnight. The second passage of recovered bacteria is treated with LL37 at two concentrations (black dash line and grey line) following the same protocol as in Fig. 1a. The second passage of bacteria exhibits similar inhibition-then-recovery dynamics of bacterial luminescent intensity, suggesting that genetic mutations do not cause the recovery in our system. See Methods Section M2. **c&d)** LL37 at 13.5μg/ml (grey line in c) leads to phosphatidylserine (PS) exposure, which has been used as a marker for bactericidal antibiotics, compared to the negative control (black line in c). Propidium iodide (PI) staining, which has been used to detect bacterial permeabilization and death, is also observed under the LL37 treatment (grey line in d), but not the negative control (black line in d). See Methods Section M3. **e)** A flow chart illustrates the experiments (Fig. 1f and 1g) that investigate the loss of Rh-LL37 (red star) activity in the presence of bacteria. Specifically, Rh-LL37 is exposed to bacterial cells for 5 hours (left). Next, medium and cells are separated using centrifugation. Fresh bacteria are inoculated into the medium portion, and the antimicrobial activity of Rh-LL37 in the medium portion is assessed using a wide-field microscope. As a control (right), Rh-LL37 is only incubated in medium without bacterial cells. See Methods Section M4. **f)** Fresh bacteria inoculated in the spent medium containing Rh-LL37 pre-exposed to bacterial cells (Fig. 1e, left) do not show any rhodamine signals inside or around the bacteria. The microscope images suggest that the Rh-LL37 loses its antimicrobial activity after pre-incubation with bacteria. Scale bar represents 10μm. See Methods Section M4. **g)** As a control (Fig. 1e, right), fresh bacterial cells demonstrate strong rhodamine intensity with Rh-LL37 pre-incubated in medium without bacteria. It implies that the Rh-LL37 retains its antimicrobial activity. Scale bar represents 10μm. See Methods Section M4.

Before investigating any collective tolerance mechanisms, we calculate the coverage of the bacterial surface by LL37 at the chosen concentrations. Specifically, we assume that 10^6^-10^7^ LL37 molecules can saturate the surface area of one *E. coli* bacterium according to a previous study using PMAP-23, because both LL37 and PMAP-23 belong to the cathelicidin family, exhibit helix conformations (Durr, Sudheendra, & Ramamoorthy, 2006; Orioni et al., 2009), and have similar estimated area-per-molecule (~550 Å^2^ for LL37 (Neville et al., 2006) and ~400 Å^2^ for PMAP-23 (Orioni et al., 2009)). We note that 6.75μg/ml LL37 corresponds to ~1.5×10^−10^ mole of LL37 molecules (M.W.=4493.3g/mol) in 100μl culture volume. Therefore, there are approximately 10^8^-10^9^ LL37 molecules to one inoculated bacterium, which is at least 10-100 fold higher than the amount of antimicrobial peptide required to saturate the surface of a single bacterium. The calculation suggests that at the sub-MIC concentration, the initial stochasticity of LL37 binding to bacterial surface is unlikely the only factor that titrates LL37 and contributes to the bacterial recovery during LL37 treatment.

Furthermore, if the amount of LL37 is not sufficient to cover the membrane areas, we expect to see an increase in the average amount of LL37 bound to bacterial membranes with a higher dose of LL37. Our flow-cytometry results counteract this argument. To track LL37, we use rhodamine labeled LL37 (Rh-LL37) that demonstrates antimicrobial activity and generates similar recovery dynamics of bacteria as unmodified LL37 (Fig. S3a) to treat wild-type *E. coli* BL21PRO (WT-BP). Previous work and our tests also show that increment of rhodamine intensity correlates with permeabilization of bacterial cytoplasmic membranes (Sochacki et al., 2011). We find that the initial distributions of rhodamine intensity in bacterial populations do not show any difference between Rh-LL37 treatments at two concentrations (one concentration with bacterial recovery; another concentration without recovery. Fig. S2c). The result implies that the average amount of Rh-LL37 bound to bacterial membranes remains the same for the Rh-LL37 treatments. The counteracting evidence between the bacterial recovery and over-coverage of LL37 molecules to bacterial membrane prompts us to investigate if there are any mechanisms that significantly reduce the effective amount of LL37 and govern the population dynamics of bacteria.

We next attempt to rule out a few canonical resistance mechanisms before investigating collective tolerance mechanisms. We first determine whether the bacterial tolerance to LL37 is heritable. Specifically, we examine whether mutations may occur in our experiments and lead to the recovery of the bacterial population. We collect bacteria (BP-lux) that recovered from LL37 treatment at 6.75μg/ml and passage them using fresh M9 medium supplemented with LL37. The passaged bacteria exhibit the same dynamics as the original bacterial populations (Fig. 1b): the passaged bacteria exhibit inhibition-then-recovery with 6.75μg/ml LL37, but no recovery with 13.5μg/ml LL37. In addition, real-time supplementation of LL37 during the recovery phase still inhibits the bacterial growth (Fig. S2d). The results indicate that the bacterial recovery is not due to random bacterial mutations or heritable resistance towards LL37.

Next, we investigate if bacteria recover due to the change of antimicrobial activity of LL37 in bacterial cultures. To track the activity and localization of LL37, we use LL37 conjugated with rhodamine. Since the conjugation of rhodamine to LL37 reduces its antimicrobial activity, we use Rh-LL37 at 54μg/ml that leads to similar dynamic as unmodified LL37 at 13.5μg/ml (Fig. S3a) to treat WT-BP and assess the antimicrobial activity of remaining Rh-LL37 in bacterial culture using microscopy. Because the results from these experiments are interpreted by comparing to negative controls, the specific concentrations we used do not impact the main conclusions of the experiments. To start, we supplement Rh-LL37 to either bacterial culture (Fig. 1e, left) or fresh medium without bacteria (Fig. 1e, right). Both samples are incubated at 37°C for 5 hours and spent medium is collected by centrifugation. To assess the antimicrobial activity of the remaining Rh-LL37 in spent medium, we inoculate fresh bacteria and monitor the co-localization of Rh-LL37 to the bacterial cells using a wide-field fluorescence microscope (Fig. 1e). Rh-LL37 that has been exposed to bacteria does not co-localize with the fresh bacteria (no detectable rhodamine intensity), indicating that the Rh-LL37 either has lost its activity or has a lower effective amount (Fig. 1f and Fig. S2e). In contrast, Rh-LL37 that has been incubated in medium without bacteria displays strong rhodamine signal around bacteria through wide-field microscopy. The results indicate that the Rh-LL37 co-localizes with the fresh bacterial cells (Fig. 1g and Fig. S2f), suggesting that the Rh-LL37 pre-incubated in medium without bacteria maintains its antimicrobial activity.

To further examine the change of antimicrobial activity of LL37 in bacterial cultures, we repeat the spent-medium experiments (Fig. 1e) with BP-lux using 13.5μg/ml of unmodified LL37. To contrast the difference between antimicrobial peptide and conventional antibiotic, we include negative controls for the AP-tolerance that are treated with 50μg/ml carbenicillin (an antibiotic that targets bacterial cell wall synthesis). Growth dynamics of fresh inoculated BP-lux in the spent medium are measured using a platereader. The working concentration of carbenicillin is determined from dosage curves where bacteria are killed within 3-4 hours (Fig. S3b). We quantify antimicrobial activity using the area between two growth curves (ABC), measured using a platereader (Fig. 2a): a higher ABC indicates more effective killing of bacteria by antibacterial agents. We find that ABC of LL37 pre-exposed to bacteria is lower than that of LL37 pre-exposed to medium without bacteria. In contrast, ABC of carbenicillin remains the same with and without pre-exposure to bacteria (Fig. 2b). Altogether, the results suggest a non-heritable mechanism that reduces LL37’s amount or activity in bacteria cultures. In the following sections, we will use his-tagged LL37 (his-LL37) and Rh-LL37 to treat *E. coli* and assess whether the free AP molecules are degraded or depleted in bacterial culture.

**Figure 2.**
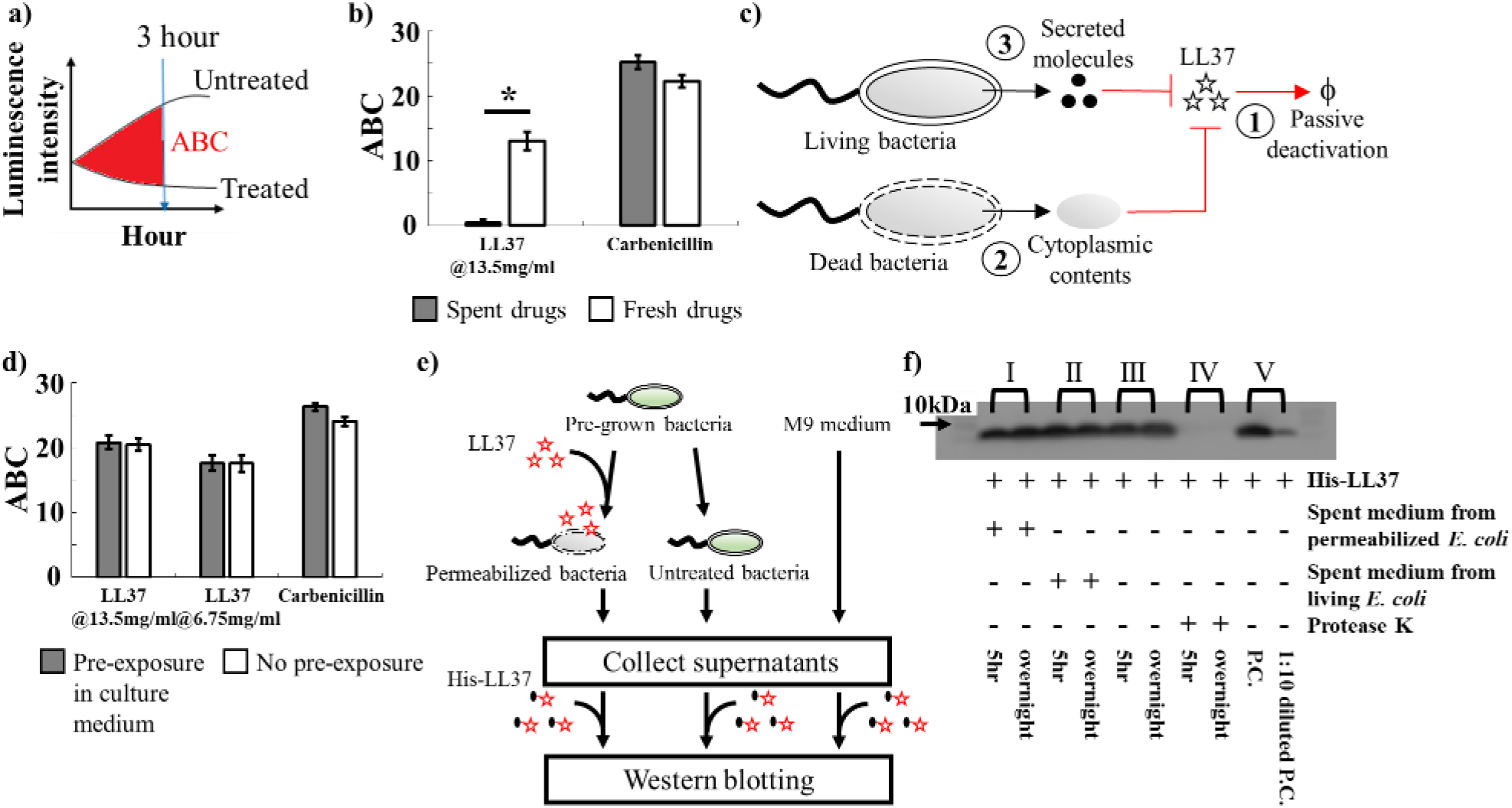
The observed bacterial population dynamics are not due to either active or passive degradation of the molecules. **a)** We define a metric named accumulated area between curves (ABC) to characterize the antimicrobial activity of drugs. It calculates the accumulated area between treated and untreated samples. Large ABC implies high antimicrobial activity. 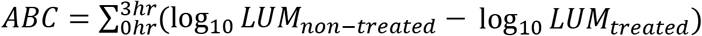. **b)** To investigate the collective tolerance mechanism, we assess the antimicrobial activity of LL37 that has been exposed to bacterial cells. Specifically, LL37 is supplemented to the bacterial culture. Next, the medium and bacterial cells are separated. The spent drug in the medium portion is collected and re-inoculated with fresh bacteria. LL37 pre-exposed to bacteria loses antimicrobial activity (left grey bar), whereas LL37 pre-exposed to culture medium without bacteria retains its antimicrobial activity (left white bar). Carbenicillin maintains its activity after pre-exposure to bacteria (right bars). Asterisk indicates significant difference (p<0.01), and error bars are standard error of the mean (SEM) from N=6. See Methods Section M4. **c)** The schematic shows possible mechanisms that may reduce the antimicrobial activity of LL37 in bacterial culture: ①. Natural degradation or self-aggregation may alter and mask functional domains of LL37. ②. Permeabilized bacterial cells may release intracellular contents that degrade or inactivate LL37. ③. Live bacteria may secrete molecules that degrade or inactivate LL37. **d)** To evaluate natural degradation or self-aggregation of LL37 in medium (①), LL37 and carbenicillin are supplemented in the medium for 3 hours before inoculation of bacteria (grey bars). Pre-incubated LL37 does not show a decrease in ABC when compared to fresh LL37. It implies that natural degradation, self-aggregation, and passive inactivation do not decrease LL37 activity within the experiment time window. Error bars are SEM from N=6. See Methods Section M5. **e)** A flow chart illustrates the experiments that investigate stability (①) of his-LL37 in medium and the degradation (② and ③) of his-LL37 in spent medium from permeabilized or live bacteria (Fig. 2f). Specifically, bacterial cells are permeabilized by LL37, and cytoplasmic contents in the spent medium are collected by centrifugation (left). Secreted molecules are collected by removing untreated bacteria from the medium (middle). As a control, medium without cells is included (right). His-LL37 is incubated in the spent medium or fresh medium and subjected to western blotting. See Methods Section M5. **f)** Quantification of the relative amount of his-tag labeled LL37 (his-LL37). We treat his-LL37 with the collected spent medium (Fig. 2e) for 5 hours or overnight to assess its degradation. The relative amount of his-LL37 is quantified using the band intensity from western blotting. His-LL37 incubated in spent medium from permeabilized bacteria (I) or live bacteria (II) does not show a reduction of band intensity. Furthermore, the relative amount of his-LL37 does not change over time in medium without bacteria, which further corroborates that LL37 does not naturally degrade in the medium (III). Proteinase K retains proteolytic activity in our reaction condition (IV). Western blotting is sensitive to a 10-fold decrease in the amount of his-LL37 (V). See replicate of western blotting results in Fig. S4a. See Methods Section M5.

### The non-heritable mechanism is not due to degradation of LL37

We study if LL37 loses antimicrobial activity through natural degradation, self-aggregation, or adhesion to culture chambers (Fig. 2c, ①). To test these possibilities, we pre-incubate LL37 for 3 hours at 37°C in the M9 medium before inoculating BP-lux and assess its antimicrobial activity by tracking bacterial luminescence using a platereader. The pre-incubated LL37 at both 6.75μg/ml and 13.5μg/ml give rise to the same ABC as fresh LL37 (Fig. 2d). As a control for AP-tolerance, both pre-incubated and fresh carbenicillin generate the same ABC. The results suggest that LL37 is not deactivated through any passive means within the time-scale of our experiments.

LL37 may be degraded by cytoplasmic contents released from permeabilized bacteria (Fig. 2c, ②). To test this hypothesis, we use western blotting to investigate if cytoplasmic contents degrade his-tagged LL37 (his-LL37). To collect cytoplasmic contents, we treat WT-BP with LL37 at 13.5μg/ml to permeabilize bacterial membranes as previously described (Fig. 1a and 1d). We then extract spent medium from permeabilized bacteria by centrifugation. The cytoplasmic contents in the spent medium directly mimic the molecular concentration and composition in the extracellular environment of a bacterial culture that has undergone LL37 treatment. The spent medium is then supplemented with his-LL37 at 37°C for 5 hours or overnight (Fig. 2e, left). We next compare the relative amount of his-LL37 incubated for 5 hours and overnight using western blotting to assess its degradation. If cytoplasmic contents degrade his-LL37, we would expect a reduced intensity of the band for 5 hours or overnight treatment compared to positive control (fresh his-LL37 at identical concentration). Western blotting does not show any difference between the band intensities of the 5 hours sample, overnight sample, and positive control (Fig. 2f-I and Fig. S4a). The result implies that the amount of his-LL37 is not reduced after either 5 hours or overnight incubation in the spent medium. We note that the western blotting is capable of distinguishing at least 10-fold decrease in the relative amount of his-LL37 (Fig. 2f-V and Fig. S4a). We repeat this experiment using whole-cell-extract (WCE) from *E. coli* BL21PRO instead of spent medium from permeabilized *E. coli*. Again, we find no degradation of his-LL37 by the WCE (Fig. S4b. SI Methods Section SI-M7). To further explore the degradation of LL37 by cytoplasmic contents, we assess the antimicrobial activity of Rh-LL37 after incubation with the spent medium from permeabilized *E. coli* using a microscope (Fig. S5a). We find that Rh-LL37 still retains its activity after pre-exposure of 5 hours to spent medium from permeabilized bacteria (Fig. S5c). Therefore, our results suggest that cytoplasmic contents released from permeabilized bacteria do not degrade LL37.

LL37 may also be degraded or sequestered by secreted molecules from live bacteria (Fig. 2c, ③). Here, we collect spent medium from WT-BP without LL37-treatment, which contains secreted molecules from bacteria. We supplement the spent medium with his-LL37 and compare its relative concentration after either 5 hours or overnight incubation to a positive control using western blotting. Again, we observe no difference between the band intensities of 5 hours sample, overnight sample and positive control (Fig. 2f-II; Fig. S4a), which implies that the amount of his-LL37 is not decreased by the spent medium. Furthermore, his-LL37 incubated in the medium without cells is not degraded between 5 hours and overnight, which corroborates that self-degradation of LL37 does not occur in our system (Fig. 2f-III; Fig. S4a). We also demonstrate that his-LL37 is still degraded by proteinase K in the reaction condition to rule out any unintended loss of protease activity in the medium (Fig. 2f-IV). Next, we explore the antimicrobial activity of Rh-LL37 after incubation with the spent medium from untreated *E. coli* using a microscope (Fig. S5a). We find that the incubated Rh-LL37 co-localizes with fresh bacterial cells, indicating that the spent medium from untreated bacterial culture does not diminish the antimicrobial activity of Rh-LL37 (Fig. S5b). Our results suggest that degradation of LL37 by secreted molecules of bacteria does not occur in our experiments.

### LL37 is absorbed by permeabilized bacteria

The above results have ruled out the loss of LL37 activity by either active or passive degradation. To shed light on the unknown mechanism that reduces LL37’s antimicrobial activity, we next investigate the mass balance of LL37 in bacterial cultures. We first incubate WT-BP with his-LL37 at a fixed concentration overnight. Unmodified LL37 is supplemented to some samples to facilitate the permeabilization of bacterial cells. Next, we collect the spent medium by centrifugation, and quantify the relative amount of remaining his-LL37 in the extracted supernatants by western blotting (Fig. 3a, left). His-LL37 supplemented in medium with no bacterial cells is quantified as a positive control (Fig. 3a, right). Indeed, the presence of bacteria (*E. coli* cells +) reduces the band intensities compared to the same condition with no cells (*E. coli* cells -. Fig. 3b-I, II, III. Fig. S6). Altogether, the findings suggest that the amount of free LL37 is reduced in the culture medium when LL37 permeabilizes bacteria, and the reduction is not due to degradation or deactivation of LL37 (Fig. 2c).

**Figure 3.**
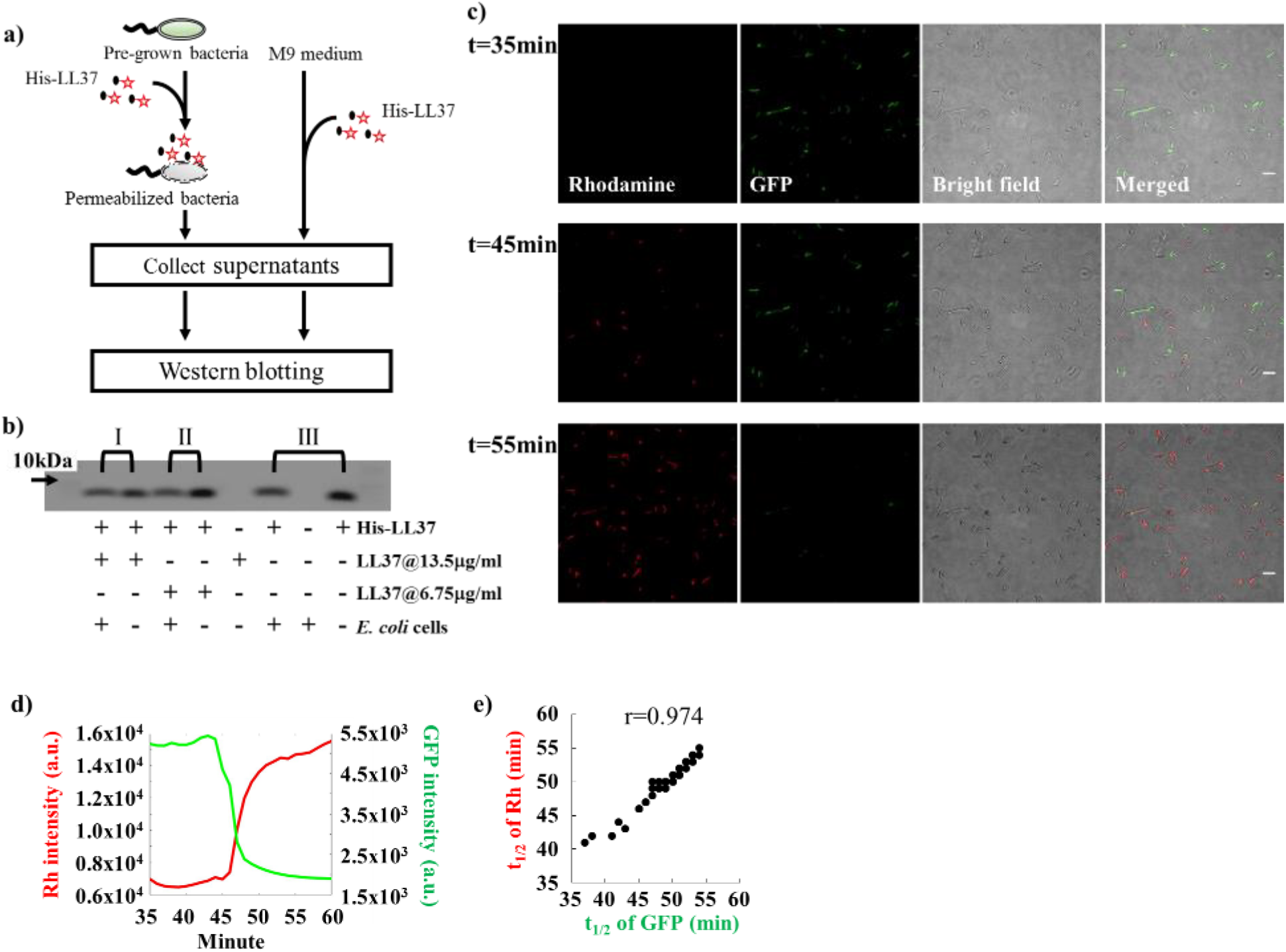
The extracellular amount of LL37 decreases in the presence of bacteria. **a)** A flow chart illustrates the experiments that investigate the conservation of mass during LL37 treatment (Fig. 3b). Bacterial culture (left) or medium without bacteria (right) is supplemented by His-LL37. Spent medium is collected, and western blotting is used to estimate the amount of remaining his-LL37 in the medium. See Methods Section M5. **b)** Western blotting shows that free his-LL37 in bacterial culture is depleted through an unknown mechanism that does not involve degradation or passive inactivation of LL37 (Fig. 2c). His-LL37 is supplemented in the bacterial culture with (I, II) and without (III) unmodified LL-37 which permeabilizes bacteria. Supernatants of the cultures are subjected to western blotting. We find that the band intensities are reduced for the samples with bacteria (*E. coli* cells +) compared to the samples without bacteria (*E. coli* cell –). See Methods Section M5. **c)** Single bacterium microscopy shows the accumulation of Rh-LL37 in bacteria. *E. coli* constitutively expressing green fluorescent proteins (GFP) is treated by Rh-LL37. Scale bar represents 10μm. See Methods Section M6. **d)** The representative dynamics of rhodamine and GFP intensity of one bacterium show that the leakage of GFP from bacterial cytoplasm correlates with the accumulation of Rh-LL37 in bacterial cells. See additional single-cell dynamics in Fig. S7a and S7b. See Methods Section M6. **e)** Single-cell GFP and rhodamine dynamics are analyzed using MATLAB to identify a correlation between the dynamics. Times to reach half of the maximum changes in GFP and rhodamine intensities (t_1/2_) are positively correlated. N=54. Methods Section M6.

To further explore the cause of free LL37 depletion in bacterial culture, we track dynamics of Rh-LL37 at the single bacterium level. *E. coli* BL21AI expresses green fluorescent proteins (BA-GFP), which are leaked outside of bacteria when their cytoplasmic membranes are permeabilized (Sochacki et al., 2011) (Fig. 3c). The bacteria are incubated in a culture chamber at room temperature for 30 minutes to allow their adhesion to the bottom surface of the chamber, and then supplemented with Rh-LL37 at 54μg/ml. At the 35th minute after the supplementation, all bacterial cells show strong GFP signals. At the 45th minute, some bacterial cells exhibit strong Rh-LL37 signals. 10 minutes later, most bacteria exhibit strong Rh-LL37 signals (Fig. 3c).

Quantification of the GFP and Rh-LL37 dynamics shows that accumulation of Rh-LL37 co-localized with bacteria coincides with the loss of cytoplasmic GFP (Fig. 3d and Fig. S7). We observe that the rhodamine intensity does not show measurable fluctuations before the drop of GFP intensity, suggesting that any binding events of Rh-LL37 (e.g., binding to the outer membrane and periplasmic space (Sochacki et al., 2011)) before cytoplasmic membrane leakage may be below the detectable limit of our wide-field microscopy. Furthermore, half-time of fluorescence signal fluctuations from bacterial cells shows a positive correlation between GFP and Rh-LL37 (with Pearson correlation coefficient (r) of 0.974, Fig. 3e). Altogether, the mass balance of LL37 and the observed LL37 absorption phenomenon suggest that LL37 permeabilizes cytoplasmic membranes of a subpopulation of bacteria, which then absorbs LL37 into their intracellular space, leading to the regrowth of the living bacteria.

To link our observations from single-cell measurements to population dynamics, we perform flow cytometry to track the fates of *E. coli* BL21PRO expressing GFP (BP-GFP) under Rh-LL37 treatment. Specifically, ~10^3^ CFU/μl of BP-GFP (See pre-growth protocol 2 in Methods Section M1) is treated with Rh-LL37 at 27μg/ml for several durations and subjected to flow cytometry. We set one threshold for GFP intensity based on the negative controls (WT-BP) that do not express GFP (Fig. 4a, and Fig. S8 for negative control). For rhodamine intensity, we first set one threshold to separate the negative control (Rh-, Fig. S8 for negative control) and Rh-LL37 associated subpopulations (Rh+). We find that the majority (~97%) of the bacterial cells has high GFP and Rh+ after 5 minutes of treatment. After 30 minutes of treatment, another subpopulation emerges at higher rhodamine intensity (Fig. 4a). We set another threshold for rhodamine intensity based on the emergent subpopulation (Rh++). At the 60th and 180th minutes, the majority of bacterial cells (~99% for both time points) has shifted to Rh++. The results strongly suggest a dynamic transition of bacterial cell states during Rh-LL37 treatment (i.e., Rh negative→①→②→③ from Fig. 4a).

**Figure 4.**
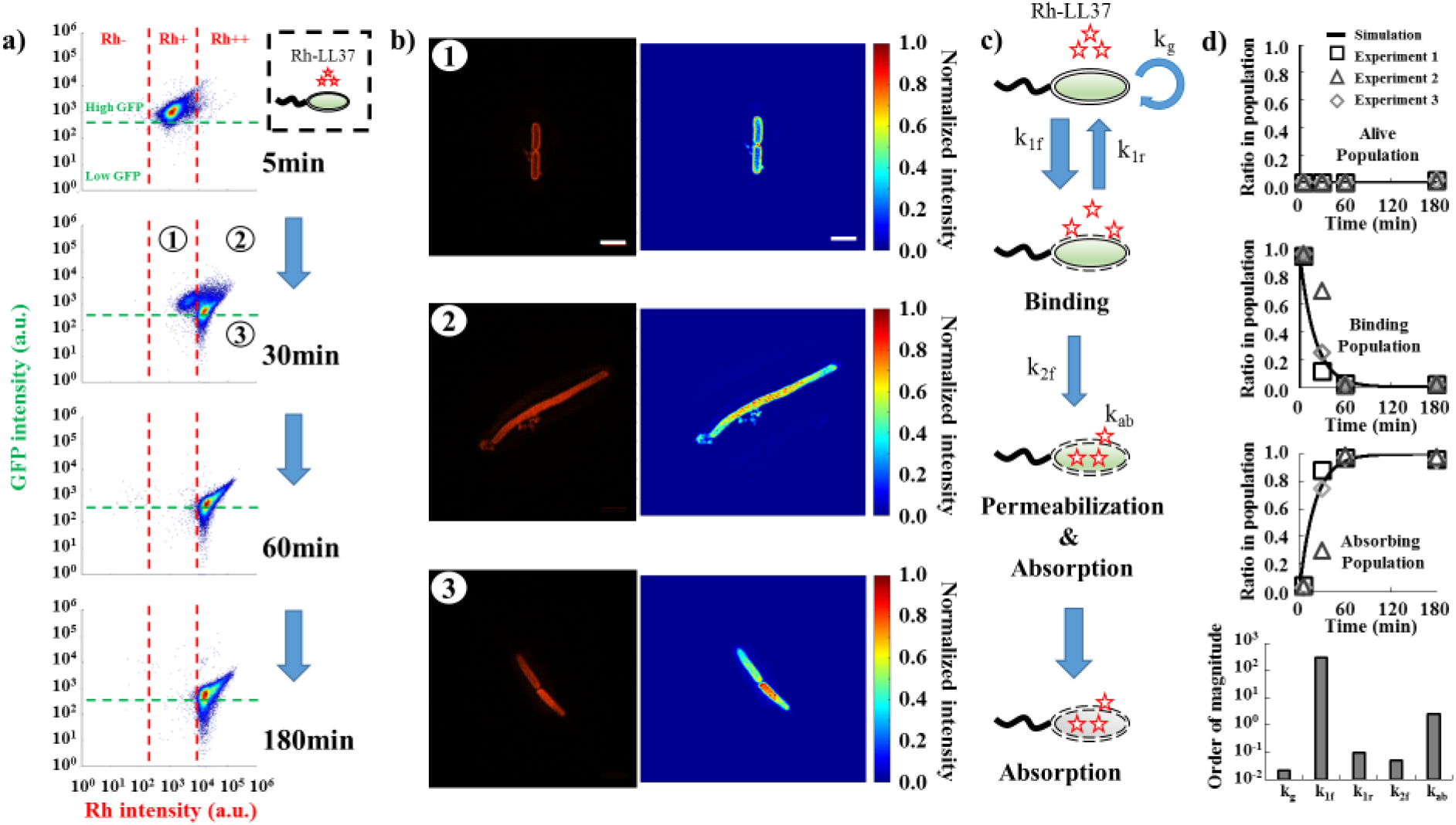
LL37 is absorbed by permeabilized bacterial cells, and the absorption causes depletion of LL37 in bacterial cultures. **a)** Flow cytometry results demonstrate the transition of bacterial states over time during Rh-LL37 treatment. *E. coli* constitutively expressing green fluorescent proteins (GFP) is treated by Rh-LL37. Green dash line separates populations with intact (high GFP) and permeabilized (low GFP) membrane. Rh negative (Rh-) represents bacterial population without Rh-LL37 association (See Fig. S8 for negative controls). Bacterial cells can transit from “high GFP, Rh positive (Rh+)” to “high GFP, Rh double positive (Rh++)” over time, and the permeabilization (transition from high GFP to low GFP) only occurs in Rh++ bacteria. See Methods Section M7. **b)** Structured illumination microscopy (SIM) reveals co-localization of Rh-LL37, bacterial membrane, and intracellular space. Bacterial populations are sorted to collect subpopulations under Rh-LL37 treatment (①, ②, and ③ from Fig. 4a). SIM images (left) and intensity heat-maps (right) show that Rh-LL37+ population (①) has Rh-LL37 co-localizes at the perimeter of the cell membrane, whereas Rh-LL37++ population (② and ③) has Rh-LL37 co-localizes at intracellular space of bacteria. See Fig. S15 for more SIM images. Scale bar represents 10μm. See Methods Section M8. **c)** Proposed model for the transition of bacterial states under Rh-LL37 treatment. We propose three bacterial states to explain the population dynamics: living population, binding population (① from Fig. 4a), and absorbing population (② and ③ from Fig. 4a). Kinetics of the transitions between states are governed by three reaction rate constants (k_1f_, k_1r_, and k_2f_). The depletion rate of Rh-LL37 from the medium by absorbing population is governed by k_ab_. Intrinsic bacterial growth has a rate constant of k_g_. See Eqn. 1 and Methods Section M9. **d)** To estimate reaction rate constants of the proposed model, we quantify ratios of each subpopulation in entire population from flow-cytometry data and estimate the kinetic parameters. Black lines represent simulation results. Black squares, grey triangles, and grey diamonds represent data from three replicates. Five reaction rate constants are estimated to be k_g_=0.022(min^−1^), k_1f_=300(min^−1^), k_1r_=0.1(min^−1^), k_2f_=0.05(min^−1^), and k_ab_=2.5(μg/ml)(min)^−1^(CFU/nl)^−1^. See Methods Section M9.

To better measure the sub-cellular localization of LL37, we treat *E. coli* BP-GFP with Rh-LL37 for 30 minutes as described above (Fig. 4a) and sort three subpopulations: high GFP, Rh+ (① in Fig. 4a), high GFP, Rh++ (② in Fig. 4a) and low GFP, Rh++ (③ in Fig. 4a). The sorted samples are subjected to high-resolution structured illumination microscopy (SIM) to identify the localization of Rh-LL37 molecules. We find that Rh-LL37 molecules accumulate at the perimeter of bacterial cells for Rh+ subpopulation (① and Fig. 4b, top), indicated by the high Rh intensities on bacterial membranes. When the bacterial cells progress to Rh++ subpopulations (② and ③), Rh-LL37 molecules co-localize with intracellular space of bacteria (Fig. 4b, middle and bottom), indicated by the higher Rh intensities in the cytoplasm than on the membranes. Furthermore, we note that bacterial population treated with Rh-LL37 at MIC demonstrates similar bacterial state transition (i.e., the transition from Rh-, Rh+ to Rh++) as the one treated at sub-MIC, implying that the AP-absorption occurs at concentrations of LL37 above MIC (Fig. S11c).

According to the observations, we propose a phenomenological model to describe the sequence of events during Rh-LL37 treatment (Fig. 4c). Specifically, we define three states of bacteria in our model: living (Rh-), binding (Rh+, ①), and absorbing (Rh++, ②, and ③) states. First, free Rh-LL37 molecules bind to bacterial cells and transfer them from “living” to “binding” state. We assume the cells can recover at a certain rate from “binding” to “living” state due to dissociation of bound Rh-LL37. The assumption is used to formulate our mathematical model (See Method Section M9). Meanwhile, bound Rh-LL37 can further progress towards permeabilizing bacterial cytoplasmic membrane (transition from “binding” to “absorbing” state). This event corresponds to the leakage of intracellular contents (reported by GFP, Fig. 3d, and 4a), as well as absorption of free Rh-LL37 molecules. Next, we build a mathematical model to quantitatively explore the proposed model (See Methods Section M9). Specifically, the progression of sequential events is governed by several reaction rate constants: kg for growth rate, k1f and k1r for forward and reverse transitions between “living” and “binding” states, k2f for transition from “binding” to “absorbing” state, and kab for AP absorbing rate (Fig. 4c). We estimate the parameters in our model by fitting to three biological replicates of the flow cytometry experiments (Fig. 4d, See Methods Section M9). The mathematical model is then extended to provide insights for population and collective tolerance dynamics of bacteria.

### The AP-absorption by permeabilized bacteria is perturbed by the presence of another bacterial strain and reduced by a peptide adjuvant

Depending on membrane surface charge, lipid composition, intracellular composition and other factors (Henzler Wildman, Lee, & Ramamoorthy, 2003; Matsuzaki, Sugishita, Fujii, & Miyajima, 1995), bacterial strains and species may display different kinetics of state transitions (k_1f_, k_1r_, k_2f_ from Fig. 4c) and AP absorption (k_ab_ from Fig. 4c, and K_ab_ which is half-maximal constant for absorption). Therefore, if the AP-absorption is true, the growth dynamics of one bacterial strain may be perturbed by the presence of the second strain during LL37 treatment (① in Fig. 5a). To start, we first estimate the kinetic parameters of bacterial state transitions and AP absorption for *E. coli* MG1655 strain that expressing GFP (MG-GFP) following the same protocol as BP-GFP (See Methods Section M7 and M9 for details). We estimate that MG-GFP demonstrates faster growth rate (larger k_g_), as well as faster recovery from “binding” to “living” state (larger k_1r_) compared to BL21PRO (Fig. 5c). Furthermore, MG-GFP also has faster permeabilization rate and absorption rate for Rh-LL37 (larger k2f and kab. Fig. 5c). The faster permeabilization and absorption rates of MG-GFP should lead to an earlier emergency of recovered subpopulation compared to BP-GFP, which is demonstrated in our flow cytometry results (Fig. S9).

**Figure 5.**
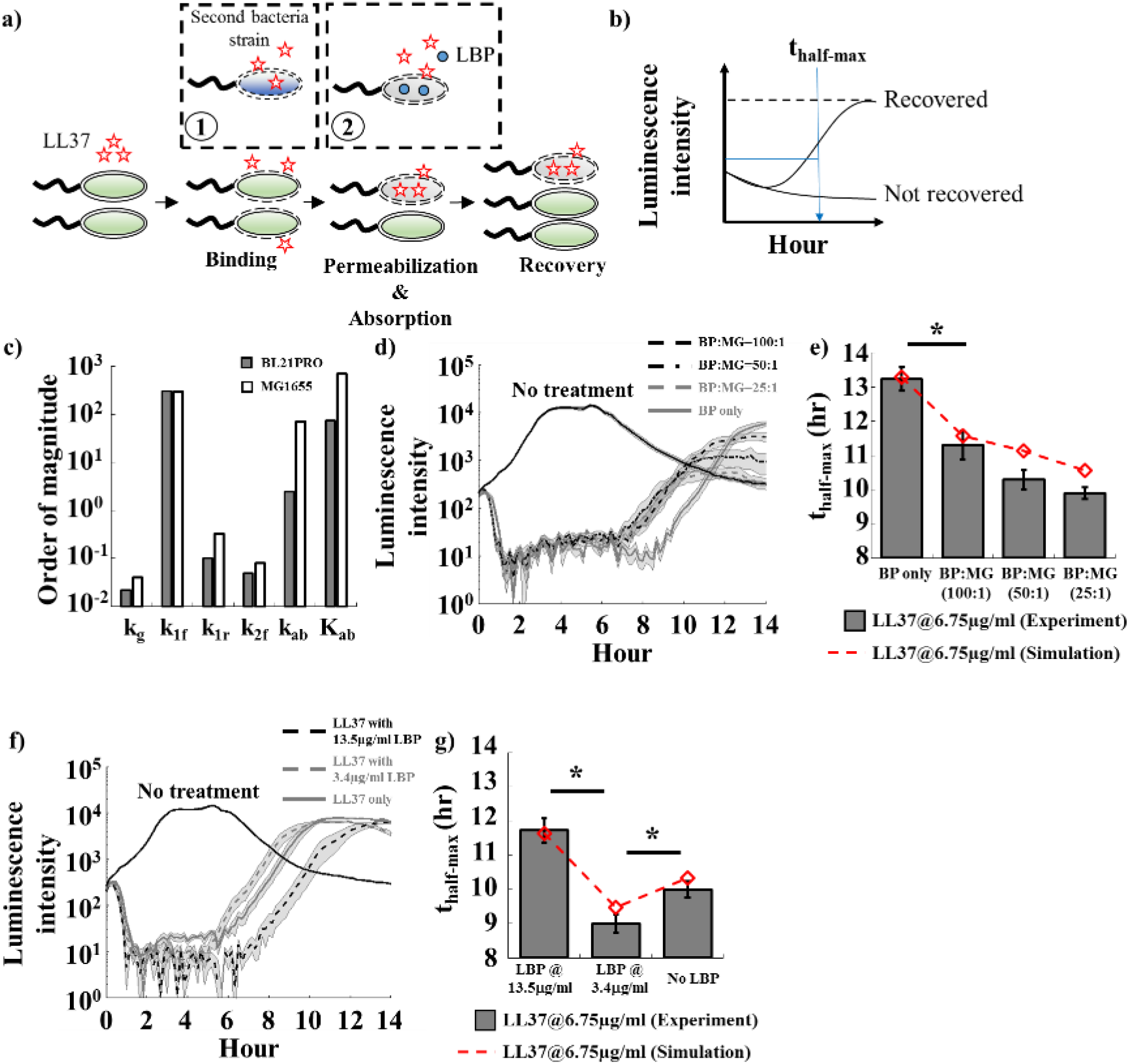
LL37 absorption by dead cells leads to cross-bacterial-strain protection and can be reduced by a peptide adjuvant. **a)** Schematic shows the perturbations of AP-absorption by another bacteria strain (①) or a competitive molecule for absorption (②). Specifically, one bacteria strain may have different LL37 absorbing kinetics (e.g., faster absorption rate) that modulate recovery of another strain during AP-treatment. In addition, a molecule (i.e., LBP) that can compete with LL37 for unintended AP-absorption may increase the antibacterial efficacy of LL37. **b)** We evaluate the recovery of the bacterial population under AP treatment using the time when the population recovers to half of its maximum growth capacity within 14 hours. **c)** Parameters estimated from flow cytometry data for BL21PRO and MG1655 are compared. See Methods Section M9. See Table S1 for the value of the parameters. **d&e)** Two strains of *E. coli* (BP-lux and MG1655) are mixed at different ratios, where total CFU of each mixture is kept constant. **d)**. Dynamic curves show the cross-bacterial-strain protection. In the presence of MG1655 (black dash and grey dash lines), the recovery times of BP-lux (which is tracked through luminescence intensity) are shifted to an earlier time compared to the one with only BP-lux (grey line). Shaded error bars are SEM from N=8. **e)**. Quantified recovery times from experiment (grey bars) and simulation (red line) showing cross-bacterial-strain protection (See SI Methods Section SI-M13 for the model, and Table S1 for estimated parameters). Asterisk indicates significant difference (p<0.01), and error bars are SEM from N=8. See Methods Section M10. **f&g)** A peptide adjuvant LBP is supplemented to the culture of BP-lux during LL37 treatment. **f)**. Dynamic curves show LBP perturbation. The recovery is delayed in the presence of LBP at 13.5μg/ ml (black dash line) but expedited in the presence of LBP at 3.4μg/ml (grey dash line) compared to the one with only BP-lux (grey line). Shaded error bars are SEM from N=8. **g)**. Quantified recovery times from experiment (grey bars) and simulation (red line) (See SI Methods Section SI-M13 for the model, and Table S1 for estimated parameters). Asterisk indicates significant difference (p<0.01), and error bars are SEM from N=8. See Methods Section M11.

Based on the difference in the kinetics, we hypothesize that the recovery of BL21PRO can be expedited in the presence of MG1655 that has a faster absorption rate (① in Fig. 5a). We first expand the mathematical model (Eqn. 1) to include MG1655 (Eqn. S1) with the estimated kinetic parameters (Fig. 5c. Table S1). In the model, two strains compete for common space (See SI Methods Section SI-M13 for model expansion). Indeed, with the same total bacterial densities, our simulation shows that recovery time of BL21PRO during LL37 treatment is accelerated by the presence of MG1655 (red dash line in Fig. 5e). To test our hypothesis, we mix BP-lux and wild-type MG1655 (WT-MG) with various CFU ratios and track the recovery of BP-lux under LL37 treatment at 6.75μg/ml using a platereader. To better quantify recovery time of BP-lux, we define a metric named thalf-max, where the population is recovered to half of its growth capacity after initial inhibition by an AP (Fig. 5b). We find that for all ratios of the two strains (BP:MG=100:1, 50:1, 25:1), the recovery of BP-lux is expedited by ~2-3 hours (Fig. 5d and 5e). Furthermore, the total initial cell density of all mixtures is tightly controlled to be identical (~10^3^ CFU/μl. See pre-growth protocol 2 in Methods Section M1), so that our observations are not affected by initial bacterial densities. The results corroborate that the AP-absorption forms a feedback response to the AP, which generates emergent collective tolerance dynamics.

LL37 is likely absorbed by several cellular components, such as intracellular DNA and lipopolysaccharide (LPS) of permeabilized bacteria due to electrostatic attraction (Bucki et al., 2007). To explore potential perturbations of LL37 absorption, we first supplement ~4ng/μl plasmid DNA extracted from *E. coli* to BP-lux culture (See pre-growth protocol 1 in Methods Section M1) under LL37 treatment. Indeed, the recovery time is reduced by ~4-5 hours with supplementation of exogenous plasmid DNA to LL37 treatment (Fig. S10a), which implies that the efficacy of LL37 is reduced. Without LL37, bacterial growth is not affected by the supplemented DNA (Fig. S10b). The results suggest that DNA is one of the intracellular components that can bind to and absorb LL37, consistent with literature data (Bucki et al., 2007).

For an AP and a bacterial strain that exhibit AP-absorption, we speculate that a peptide can compete with the absorption of the AP by intracellular components, and delay the depletion of free AP molecules. We use an LPS-binding peptide (LBP) that has been shown to prevent physical interaction between LL37 and LPS (Bucki et al., 2007). Consistent with our expectation, we find that LBP at 13.5μg/ml delays recovery time of BP-lux (See pre-growth protocol 1 in Methods Section M1) by ~2 hours compared to no LBP culture under LL37 treatment (Fig. 5f). We note that LBP alone does not inhibit bacterial growth at the tested concentrations (Fig. S11a). However, LBP displays a concentration-dependent effect. That is, LBP delays bacterial recovery at high concentration (13.5μg/ml) but expedites it at low concentration (3.4μg/ml) (Fig. 5f and 5g). Flow cytometry results show that LBP at 13.5μg/ml delays the transition from “binding” to “absorbing” state (Fig. S11b). The results suggest that LBP can inhibit both membrane-permeabilization and intracellular absorption of LL37 molecules, which leads to the concentration-dependency effect. We expand the mathematical model (Eqn. 1) to implement the proposed actions of LBP (Eqn. S2), and the simulation results agree with our experiments (red dash line in Fig. 5g, and SI Methods Section SI-M13 for model expansion). The results suggest that peptide adjuvants may be supplemented to APs to reduce or abolish AP-absorption and enhance the efficacy of APs. Further work using machine learning approaches may be used to improve the efficacy of the peptide adjuvants in abolishing AP-absorption (Lee, Fulan, Wong, & Ferguson, 2016; Lee, Wong, & Ferguson, 2017).

## Discussion

Through a series of deductive experiments, we discover a bacterial collective tolerance mechanism towards AP, in which LL37 permeabilizes and kills a subpopulation of *E. coli*, which then absorbs the LL37 into its intracellular space, leading to regrowth of living bacteria. The collective tolerance can only occur at the population level because the permeabilized and dead bacteria, in turn, absorb LL37, enhancing the escape of other bacteria in the same population. We rule out classical resistance mechanisms of bacteria, including bacterial mutation and proteolytic cleavage of LL37 (Fig. 1 and 2). We also show that short half-life and passive inactivation of LL37 are not the underlying mechanisms of the LL37-tolerance in our system (Fig. 2d and 2f-III). Both flow cytometry and single-cell microscopy corroborate the role of AP-absorption, as well as suggest a phenomenological model for bacterial population dynamics during treatment (Fig. 3 and Fig. 4). Furthermore, permeabilized bacteria absorb LL37 at both sub-MIC and MIC concentrations. Facilitated by a mathematical model, we demonstrate cross-bacterial-strain protection of bacteria against LL37 due to AP-absorption, as well as a potential peptide adjuvant to tackle the tolerance mechanism (Fig. 5).

Furthermore, our findings show that the AP-absorption is a major process that influences bacterial population dynamics during AP-treatment. For example, if the bacterial recovery (Fig. 1a) is merely due to the growth of some lucky cells that are not bound by AP, the recovery time should be independent of the presence of a second bacterial strain in the population with a constant initial-bacterial-density (Fig. 5). Instead, the observed cross-bacterial-strain protection against LL37 suggests the critical role of AP-absorption in regulating population dynamic during AP treatment. Furthermore, if the binding of AP to live bacterial membrane is the only factor that controls population dynamics, the supplementation of LBP should always expedite bacterial recovery because of competitive binding between LBP and LL37 to bacterial membrane. Instead, we find that LBP at 13.5μg/ml delays the bacterial recovery, which highlights the critical role of AP-absorption in governing the population dynamics.

The AP-absorption tolerance mechanism may be generalizable to other bacterial species and APs for a few reasons. First, the absorption of APs likely relies on generic electrostatic interactions between APs and bacterial components, which are ubiquitous across bacterial species. However, the kinetic of events leading to AP-absorption likely depends on the composition of membranes and cytoplasms that control the insertion and transport of antimicrobial peptides. Second, bacterial permeabilization is a common mechanism-of-action of a major class of APs. Upon the initial permeabilization, cationic APs may diffuse into cells and bind to negatively-charged cellular components. Since the electrostatic interaction is not unique to LL37, other cationic APs are likely to be tolerated by bacteria through the same absorption mechanism. Indeed, we may investigate the AP-absorption mechanism using similar recovery dynamics of *E. coli* under the treatment of indolicidin and bac2A (APs originated and derived from bovine neutrophils) (Fig. S12a and S12b).

The discovery of a novel collective tolerance mechanism based on AP-absorption by dead bacteria spawns a new research area with several open questions. From a qualitative point of view, intrinsic heterogeneity of bacterial population may cause the stochastic bifurcation of cell states during treatment (e.g., some bacterial cells are permeabilized faster than others due to the intrinsic heterogeneity). It is unclear if any genes or proteins are associated with the heterogeneous behavior during AP treatment, which may be investigated through cell-sorting and mRNA profiling. In addition, we have used LBP to reduce the AP-absorption and improve the efficacy of LL37. APs may be sequestered by multiple negatively-charged bacterial components including DNA, peptidoglycan (Sochacki et al., 2011), lipopolysaccharide, and f-actin (Bucki et al., 2007). Designing adjuvant molecules that compete for AP-absorption may provide a new way to improve AP efficacy. From a quantitative point of view, AP-absorption is a highly dynamic process that has the potential to generate emergent dynamics. The kinetics of bacterial death, AP binding, bacterial recovery from bound AP affect dynamics of AP absorption, which in turn affect population dynamics under AP treatment. Our study also reveals a negative feedback loop between an AP and bacteria. Specifically, an AP permeabilizes bacteria and induce bacterial cell death, but the dead bacterial cells, in turn, absorb AP and diminish the efficacy of the treatment. Different from previous studies on collective AP-tolerance, the feedback loop highlights the role of bacterial death on population survival during AP treatment, which may suggest a new direction towards improving AP efficacy by perturbing the feedback loop. Furthermore, we have shown that even the same species (*E. coli*), but different strains (BL21PRO and MG1655) demonstrate different kinetics for AP-absorption, which are sufficient to generate the cross-bacterial-strain protection against LL37. It remains unclear how APs could dynamically shape the composition of a multi-species/strains environment under treatment due to AP-absorption, especially when interactions between species/strains are involved.

## Acknowledgements

We appreciate the discussion of the manuscript with members of Tan lab. This work is supported by Society-in-Science: Branco-Weiss Fellowship to C.T. Cell-sorting is supported by the National Institutes of Health under award number S10OD018223.

## Author contributions

F.W. and C.T. designed the experiments. F.W. performed the experiments and analyzed the results. All authors wrote the manuscript.

## Materials and Methods

### Bacterial strains and chemicals (M1)

*Escherichia coli* BL21PRO strain carrying plasmid that constitutively expresses *lux* genes (BP-lux) was used for measurement in a platereader. Wild-type *E. coli* BL21PRO (WT-BP) and *E. coli* BL21AI constitutively expressing GFP (BA-GFP) were used for tracking Rh-LL37 dynamics with wide-field microscopy. *E. coli* BL21PRO and MG1655 expressing GFP (BP-GFP and MG-GFP) were used for flow cytometry and structured illumination microscopy (SIM). All strains were maintained as glycerol stocks at −80°C for long-term storage or on LB agar plates at 4°C for short-term storage. Bacteria were grown overnight at 37°C in Luria Broth (LB) (VWR) before experiments. Two pre-growth protocols were used before any treatments to ensure bacteria cells enter exponential growth phase:

#### Pre-growth protocol 1

Fresh overnight cultures were diluted 1:1,000 into the M9 minimal medium (VWR) supplemented with 0.2% glucose and 0.2% casamino acids without any antibiotic selection and grown at 37°C on a shaker for 2 hours.

#### Pre-growth protocol 2

Fresh overnight cultures were diluted 1:1,000 into the M9 medium without any antibiotic selection. The cultures were grown at 37°C on a shaker for 3 hours. To better control bacterial cell density, OD600, luminescence or GFP intensity of the pre-grown cultures was measured, and its CFU/μl was back-calculated according to calibration curves. The pre-grown cultures were then diluted with M9 to have ~10^3^ CFU/μl for later experiments. CFU was also performed for diluted cultures to check the quality of density control.

Unmodified LL37 was purchased from AnaSpec. Rhodamine-conjugated LL37 (Rh-LL37) was purchased from Rockland. The APs were reconstituted in nanopore water (Thermo Scientific) before use and stored at −20°C. Six histidine residues were fused to N-terminus of LL37 (SI Methods Section SI-M4 and Fig. S13). The fused peptide was expressed from a high copy number plasmid (pET15bL) using BL21DE3. Specifically, 500μl of fresh overnight BL21DE3/pET15bL-his-LL37 culture was inoculated into 200ml LB medium and incubated at 37°C with 200rpm shaking until it reached exponential growth phase. Next, IPTG was supplemented at 0.4mM working concentration to induce the expression of his-LL37 for 3 hours. Bacteria were then harvested using centrifugation (10,000 g, 10 minutes). Each gram (wet weight) of cell pellets was re-suspended with 5ml of a binding buffer (200mM NaCl and 25mM Tris-HCl in water). We then lysed bacteria through sonication (QSonica Q125, 67% amplitude; 8 cycles of 15 seconds “ON” and 45 seconds “OFF”), and collected cytoplasmic contents through high-speed centrifugation (25,000 g, 1 hour). His-tag labeled LL37 was purified with nickel column (Fisher Scientific) and stored at −20°C for future experiments (SI Methods Section SI-M5). Carbenicillin was purchased from Sigma.

LBP was synthesized from Biomatik according to the amino acid sequence obtained from the previous literature (Araña et al., 2003): RVQGRWKVRKSFFK with FITC linked to N-terminus. The LBP was reconstituted in nanopore water before use and stored at −20°C.

### Measurement of bacterial growth dynamics using a platereader (M2)

BP-lux was grown with pre-growth protocol 1. 100μl of bacterial culture was aliquoted into each well of a black flat bottom 96-well microplates (Corning Costar). LL37 was supplemented to the wells at 6.75μg/ml or 13.5μg/ml working concentrations (Fig. 1a). Time series of luminescence was measured using Tecan M1000Pro platereader at 37°C with shaking (orbital, 20s every min). Parameters for luminescence measurement were automatic attenuation and 1,000ms integration time.

To test for bacterial mutation (Fig. 1b), BP-lux was treated with LL37 at 6.75μg/ml, and luminescence was tracked in the platereader as described above. When a bacterial population started to recover (~7 hours after treatment), 50μl of the bacterial culture was extracted from the wells and added into 3ml of LB medium without any antibiotic selection to initiate a new overnight culture. The new overnight culture was then treated with LL37 again following the same protocol as stated above.

### Phosphatidylserine (PS) exposure, propidium iodide (PI) staining, and flow cytometry (M3)

Annexin V-FITC Apoptosis detection kit (Sigma) was used to measure PS exposure and PI straining. Specifically, WT-BP was grown using pre-growth protocol 1. LL37 was added to the culture at 13.5μg/ml working concentration. After 2 hours of LL37 treatment, 1ml of culture was collected and centrifuged at 10,000 g for 10 minutes. Cell pellets were re-suspended with Annexin binding buffer provided in the kit. Annexin V and PI dyes were added as described in the manual, and samples were incubated in the dark for 30 minutes at room temperature. Stained samples were diluted 1:50 into PBS and flow cytometry was performed using FACScan 5-color cytometer. Parameter settings of the flow cytometer were: Lasers: 488nm blue and 640nm red; Detectors: 530-580nm FITC and 627-666nm PI; Voltages: 295 SSC, 551 FITC, and 458 PI. FCS-SSC gate was created for bacterial cells based on bacteria under no treatment. Around 10,000 events within FCS-SSC gate for bacterial cells were collected for each sample.

### Testing of LL37 and carbenicillin in spent medium (M4)

For the platereader assays (Fig. 2b), we treated BP-lux (pre-growth protocol 1) with either LL37 or carbenicillin and incubated them in 96-well plates. After 3 hours of incubation at 37°C, treated cultures were collected in 1.5ml Eppendorf tubes and centrifuged at 10,000 g for 10 minutes to collect the supernatants. Extracted supernatants were added to 96-well plates, and inoculated using the fresh overnight culture of BP-lux at 1:50 dilution ratio. Luminescence was measured in the platereader. Negative controls contained only M9 medium with LL37 or carbenicillin and were subjected to the same procedure as stated above.

For the microscopy assay (Fig. 1e-g), WT-BP (pre-growth protocol 1) was treated with Rh-LL37 at 54μg/ml in 96-well plates. After 5 hours of incubation at 37°C, treated cultures were collected in 1.5ml tubes and centrifuged at 10,000 g for 10 minutes. 10μl of pre-grown bacteria (pre-growth protocol 1) were relocated in microscopic slide chamber (μ-Slide Angiogenesis, Ibidi) following the supplementation of 20μl of the supernatants. The chamber was incubated at room temperature for 2 hours. Images were acquired using Nikon Eclipse Ti microscope with 100x objective (Technical Instruments, CA). The settings for microscope filters were: 450-490nm excitation and 500-550nm emission for GFP; 532-557nm excitation and 570-640nm emission for rhodamine. The exposure times for both GFP and rhodamine were 100ms.

### Platereader and western blotting to study degradation of LL37 (M5)

To test the self-deactivation of LL37 (Fig. 2d), we supplemented LL37 or carbenicillin in M9 medium without bacteria and incubated them for 3 hours at 37°C on a shaker. The fresh overnight culture of BP-lux bacteria was then inoculated at 1:50 dilution ratio into the M9 medium with either pre-incubated or fresh drug molecules. Population dynamics of the bacteria were tracked using the platereader.

To examine degradation of LL37 (Fig. 2e and 2f), we treated WT-BP (pre-growth protocol 1) with or without LL37 at 13.5μg/ml in 96-well plates at 37°C for 4 hours. The supernatants were then collected by centrifugation (25,000 g, 1 hour). Next, we mixed the collected supernatants with purified his-LL37 at 1:1 volumetric ratio in PCR tubes. The controls contained M9 medium, purified his-LL37, or protease K at 1mg/ml (Thermo Scientific). The samples were incubated at 37°C for either 5 hours or overnight and subjected to western blotting with the His-tag antibody (Thermo Scientific). Specifically, samples were run through Mini-PROTEAN TGX Precast Gel (Bio-Rad) and transferred to nitrocellulose membranes using Trans-Blot Turbo RTA Nitrocellulose Transfer kit (Bio-Rad). The transferred membranes were blocked using 5% milk (Biotium) in TBST (1x TBS and 0.1% Tween 20 in water) for 1 hour.

Solutions for western blotting were prepared as follow: primary antibody (6x-His Epitope Tag Antibody from mouse, Thermo Scientific): 1:3,000 dilution in 3% BSA (in TBST); secondary antibody (Goat anti-Mouse IgG Secondary Antibody, HRP conjugate, Thermo Scientific): 1:20,000 dilution in 3% BSA (in TBST); washing buffer: 0.2% milk in TBST. The staining process was performed as follow: incubated in primary antibody for 1.5 hours → washed three times in washing buffer for 10 minutes each → incubated in secondary antibody for 1 hour → washed three times in washing buffer for 10 minutes each. Last, HRP on membranes was detected using Clarity Western ECL Blotting Substrates (Bio-Rad) and PXi gel imager (Syngene). The incubation and washing for western blotting were all performed at room temperature on an orbital horizontal shaker. Positive controls (P.C.) were prepared by mixing his-LL37 and M9 medium at 1:1 volumetric ratio and subjected to western blotting.

To investigate depletion of free LL37 in medium (Fig. 3a and 3b), we mixed 10μl of purified his-LL37 to 10μl of WT-BP (pre-growth protocol 1). Next, we supplemented LL37 at either 6.75μg/ml or 13.5μg/ml to permeabilize bacteria. The mixtures were incubated at 37°C overnight in PCR tubes. Supernatants of the cultures were extracted using centrifugation (25,000 g, 1 hour). The negative control contained M9 medium and his-LL37. The collected supernatants were subjected to western blotting as described above.

### Tracking dynamics of Rh-LL37 through wide-field fluorescence microscopy (M6)

BA-GFP was pre-grown using protocol 1 supplemented with 0.2% arabinose to induce expression of GFP. We aliquoted 30μl of bacterial culture to a slide chamber and allowed the cells to settle down for 30 minutes at room temperature. Next, we added Rh-LL37 at 54μg/ml working concentration to the chamber. Images were recorded every 1 minute with the 100x objective for 1 hour. The microscope settings were the same as stated above. Microscope images were analyzed using ImageJ. Specifically, bacterial cells that were visualized throughout all time points were selected. Integrated intensities of rhodamine and GFP at different time points were quantified for each selected bacterium and input into MATLAB to generate Figure 3d&e.

### Tracking transitions of bacteria states during Rh-LL37 treatment using flow cytometry (M7)

BP-GFP and MG-GFP were pre-grown following pre-growth protocol 2 supplemented with 0.4mM IPTG to induce expression of GFP. The initial cell density for this experiment was well controlled so that we could compare quantitative results across two strains. 100μl of pre-grown culture was aliquoted in 96-well plate, and Rh-LL37 was supplemented at 27μg/ml. The samples were incubated in platereader at 37°C with same shaking protocol as described before. At specific time points, the samples were added to 1ml of 4% PFA and stored on ice before flow cytometry. Bacteria with no Rh-LL37 treatment was included as a control to gate-out noise signal based on FSC and SSC. WT-BP and WT-MG were included for negative controls of GFP intensity. Flow cytometry was performed on Thermo Fisher Attune NxT flow cytometer. At least 20,000 events within FSC-SSC gate for bacterial cells were collected.

### Bacteria cell-sorting and structured illumination microscopy (SIM) (M8)

BP-GFP treated with Rh-LL37 at 27μg/ml was prepared as described for the flow cytometry experiment. Samples were collected after 30min of treatment and diluted in 4% PFA. Cell sorting was performed using Beckman Coulter MoFlo Astrios Cell Sorter at UC Davis Flow cytometry Core Facility. BP-GFP with no treatment was run through cell sorter first to gate-out noise based on FSC and SSC and create thresholds for GFP negative and Rh negative. Next, the threshold to separate Rh+ and Rh++ was set based on grouping of subpopulations (See Fig. 4a for example). Bacterial cells from three representative regions (①, ②, and ③ in Fig. 4a) were sorted and stored on ice before imaging using SIM.

To prepare samples for SIM, 100μl of 2% agarose was melt and dropped on a glass slide. A cover glass was placed on the top of agarose to flat the surface until it was dry. 10μl of the sorted sample was dropped on the top of agarose and mounted with a cover glass. SIM was performed using Nikon Structured Illumination “Super-Resolution” microscope equipped with 488nm, 565nm laser lines and 100x objective at UC Davis Microscopy Imaging Facility. Raw images were acquired with 3D-SIM mode and reconstructed using provided software (Nikon Elements). Reconstructed images were analyzed using ImageJ (Fig. 4b, left) and MATLAB (Fig. 4b, right). To obtain the heat-map of Rh-LL37 localization (Fig. 4b, right), each image was imported to MATLAB and normalized with the highest intensity in the image.

### Mathematical model and parameter estimation using flow cytometry data (M9)

We construed a deterministic model using a system of ordinary differential equations (ODE) to explore the population dynamics of single bacterial strain under LL37 treatment (Eqn. 1).

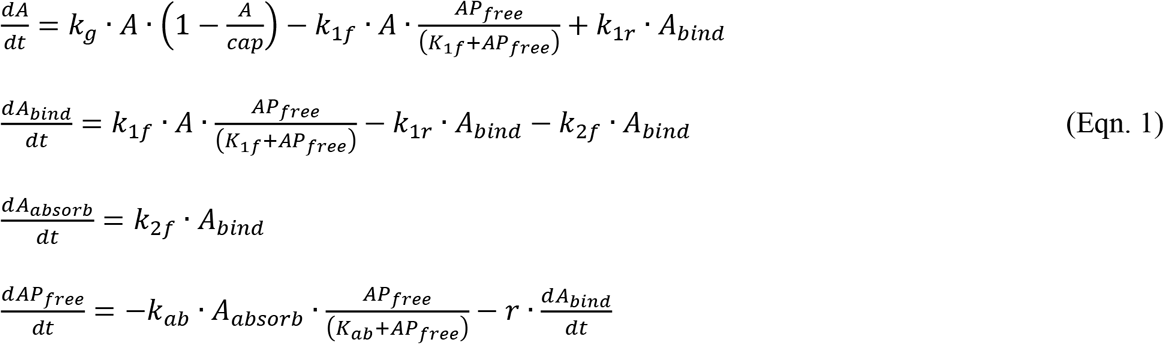

Here, A represented bacteria that were not affected by LL37 (living population); A_bind_ represented bacteria that had LL37 bound to their perimeters (binding population); A_absorb_ represented bacteria that had been permeabilized and were absorbing free LL37 molecules (absorbing population). AP_free_ was free LL37 molecules in the medium. Five reaction rate constants were kg, k_1f_, k_1r_, k_2f_ with a unit of [min]^−1^, and k_ab_ with a unit of [μg/ml][min]^−1^ [CFU/nl]^−1^. K_1f_ and K_ab_ were half-maximum constants with a unit of [μg/ml]. r was the proportional coefficient between A_bind_ and bound LL37 molecules with a unit of [μg/ml][CFU/nl]^−1^. We set r=0.05, which corresponded to roughly 6.7×10^6^ LL37 molecules to one bacterium, based on the saturation level of the AP PMAP-23 on the bacterial membrane (Roversi et al., 2014). K1f was approximated to be 45μg/ml (roughly equal to 10μM) using apparent dissociation constant of LL37 on bio-membrane (Sood, Domanov, Pietiainen, Kontinen, & Kinnunen, 2008). The growth rate of A was governed by kg, and had a capacity of cap=100 (corresponding to ~100 fold changes from initial density to growth capacity from Fig. 1a). kg was estimated to be 0.022 for BL21PRO or 0.04 for MG1655 so that the culture took ~6-7 hours for BL21PRO or ~3-4 hours for MG1655 to reach its capacity, which was similar to our experimental results with pre-growth protocol 2 (Fig. S14). Forward and reverse transitions between A and Abind were represented using 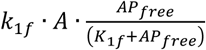 and *k*_1r_∙A_bind_ respectively. Permeabilization of Abind was governed by k_2f_. AP_free_ could be depleted from medium through either the absorption 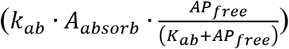 or binding to bacterial membrane 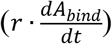. The depletion of free LL37 molecules through binding was directly proportional to A_bind_.

To estimate kinetic parameters, we first extracted ratio of each subpopulation from flow cytometry data of BP-GFP and MG-GFP under Rh-LL37 treatment. Specifically, two thresholds for rhodamine intensity were set as described for samples collected at different time points. Ratios of Rh-, Rh+, and Rh++ to entire population were recorded and used as experimental data for parameter estimation. We used MATLAB function *fmincon* to obtain the first estimation based on our mathematical model (Eqn. 1) using three replicates of flow-cytometry data. The loss function for *fmincon* calculated the summation of the square difference between simulated and three replicates of experimental data. Next, parameters were further refined to fit both the flow cytometry and recovery time measurements (See Table S1 for simulation parameters).

### Tracking recovery of BP-lux in the presence of WT-MG (M10)

BP-lux and WT-MG were pre-grown following pre-growth protocol 2 to ensure tightly controlled initial density. Then, two pre-grown cultures were mixed with various volumetric ratios to create BP:MG=100:1, 50:1, and 25:1 mixtures. 100μl of each mixture was aliquoted in 96-well plate and supplemented with LL37 at 6.75μg/ml. Recovery dynamics of BP-lux were tracked through luminescence using the platereader.

### Perturbation of AP-absorption by LBP (M11)

BP-lux was pre-grown following pre-growth protocol 1. 100ul of pre-grown BP-lux was aliquoted in 96-well plate, and LBP was supplemented at designed concentrations. LL37 was then added at 6.75μg/ml. Recovery dynamics of BP-lux were again tracked through luminescence using the platereader.

### Statistical test (M12)

All statistical tests were performed using at least six replicates. To compare the means of two groups, a one-tail t-test was used with p<0.01. Pearson correlation coefficient was calculated to estimate the linear correlation between two variables.

